# TaxMapper: An Analysis Tool, Reference Database and Workow for Metatranscriptome Analysis of Eukaryotic Microorganisms

**DOI:** 10.1101/174227

**Authors:** Daniela Beisser, Nadine Graupner, Lars Grossmann, Henning Timm, Jens Boenigk, Sven Rahmann

## Abstract

**Background:** High-throughput sequencing (HTS) technologies are increasingly applied to analyse complex microbial ecosystems by mRNA sequencing of whole communities, also known as metatranscriptome sequencing. This approach is at the moment largely limited to prokaryotic communities and communities of few eukaryotic species with sequenced genomes. For eukaryotes the analysis is hindered mainly by a low and fragmented coverage of the reference databases to infer the community composition, but also by lack of automated workflows for the task.

**Results:** From the databases of the National Center for Biotechnology Information and Marine Microbial Eukaryote Transcriptome Sequencing Project, 142 references were selected in such a way that the taxa represent the main lineages within each of the seven supergroups of eukaryotes and possess predominantly complete transcriptomes or genomes. From these references, we created an annotated microeukaryotic reference database. We developed a tool called TaxMapper for a reliably mapping of sequencing reads against this database and filtering of unreliable assignments. For filtering, a classifier was trained and tested on sequences in the database, sequences of related taxa to those in the database and randomly generated sequences. Additionally, TaxMapper is part of a metatranscriptomic Snakemake workflow developed to perform quality assessment, functional and taxonomic annotation and (multivariate) statistical analysis including environmental data. The workflow is provided and described in detail to empower researchers to easily apply it for metatranscriptome analysis of any environmental sample.

**Conclusions:** TaxMapper shows superior performance compared to standard approaches, resulting in a higher number of true positive taxonomic assignments. Both the TaxMapper tool and the workflow are available as open-source code at Bitbucket under the MIT license: https://bitbucket.org/dbeisser/taxmapper and as a Bioconda package: https://bioconda.github.io/recipes/taxmapper/README.html.

## Background

### Motivation and goals

Metatranscriptome sequencing of diverse ecosystems is becoming a common methodology in many research institutions, and large scale sampling campaigns such as the Marine Microbial Eukaryote Transcriptome Sequencing Project (MMETSP, [1]) and the Tara Oceans expedition [2] have contributed to a growing amount of available environmental sequencing data. However, the analysis of the resulting short read sequences is still far from routine, especially for unicellular eukaryotic organisms, due to what was termed by Escobar-Zepeda *et al.* as “the neglected world of eukaryotes in metagenomics” [3]. This is particularly severe since microscopic eukaryotes (protists) constitute a paraphyletic taxon [4] spread over the whole eukaryotic tree of life and represent the bulk of most major groups, whereas multicellular lineages are confined to small corners [5]. Protists occur at high abundance in almost all habitats, e.g. in freshwaters, oceans, biofilms and soils [5, 6, 7, 2, 8, 9]. They maintain ecosystem functions, as they are responsible for most planktonic primary production [10], are the most important feeders of bacteria [11, 7] and key players in the regulation of element cycling, particularly carbon [7, 12].

Perhaps surprisingly then, protists are poorly covered by genomic reference databases despite their broad diversity, and if at all, only few model species are present. Therefore, most recent metatranscriptome approaches were designed for prokaryotes, which offer more complete databases (e.g. NCBI) in contrast to eukaryotes. Here, efficient mapping approaches, such as BWA or Bowtie, and methodologies allowing few differences to the reference sequences (e.g. k-mer indices) can be used. It is frequently possible to obtain taxonomic assignments even down to species level.

In contrast, few genome sequences from eukaryotes exist, and those that do are not well balanced across the main lineages of the eukaryotic tree of life, and therefore do not reflect the diversity within these lineages. The main focus of publicly available genomes lies on the Opisthokonta (Fungi/Metazoa group), including many animals, in particular model organisms, and Viridiplantae (green plants, containing Streptophyta and Chlorophyta) with an emphasis on crop plants. For example, in the NCBI database the available genomes in these two groups already represent 96% of the available genomes for eukaryotes, whereas eukaryotic genomes represent 43% of all genomes from the three domains (bacteria: 54%, archaea: 3%, NCBI June 2017).

The diversity of microbial eukaryotes is strongly underrepresented and database searches that aim at an assignment of metatranscriptomic reads on species level will, for the most part, be incorrect. This is caused by the fact that neither the species nor a close relative are included in the database and by the disproportional coverage of taxonomic groups leading to misassignments of reads to incorrect taxa by chance. In addition, available databases are often too large to be used in their entirety to map or search with millions of metatranscriptomic sequences on the read level.

A possible way out (taken here) is to restrict the taxonomic assignment to broader taxonomic groups, using appropriate reference organisms for each group. In turn, this requires a different approach to the similarity search, allowing to find more distantly related sequences. Since such similarity search tools are more time consuming, a reasonable search time can only be obtained by restricting the analysis to smaller reference databases.

Many existing approaches base their taxonomic assignments on selected sequenced marker genes. However, for a joint taxonomic and functional analysis (which taxonomic group performs which functions?), it is necessary to assign *each single read* to a taxonomic group and to a protein family.

Our goal was therefore to design, test and provide a comprehensive tool and workflow for eukaryotic metatranscriptome analysis, encompassing everything from preprocessing to integration of environmental data. A large impediment, as already mentioned, was a missing reference for the taxonomic assignment of sequences, which we constructed for all major taxonomic groups based on 142 publicly available transcriptomes and genomes. Our tool TaxMapper assigns taxonomic information to each read by mapping to the database using a reduced amino acid alphabet, and subsequently filtering of unreliable assignments. It is part of an automated rule-based Snakemake workflow developed to perform quality assessment and both functional and taxonomic annotation, as well as (multivariate) statistical analysis including environmental data.

In this work, we (i) describe the microeukaryotic reference database, (ii) present the TaxMapper software for taxonomic mapping and filtering of reads, and (iii) provide a detailed step-wise instruction on how to analyse metatranscriptomes from eukaryotic microorganisms using a modular workflow.

### Related work

#### Metatranscriptome workflows

Existing metatranscriptome workflows often focus on bacterial composition, like Leimena *et al.* [13] who describe in detail an analysis pipeline for prokaryotic datasets. Other studies construct pipelines for subparts of the analysis, including Goncalves *et al.* [14] who constructed an R-based pipeline for pre-processing, quality assessment and expression estimation of RNA sequence datasets, and Marchetti *et al.* [15] who provide an R package for differential expression analysis of metatranscriptome sequences starting from a count matrix of genes and a phylogenetic annotation. For our purposes, these approaches have two disadvantages: (i) they provide no complete executable workflow, and (ii) the available workflow parts cannot be easily adapted to eukaryotic data.

#### Metatranscriptome analysis tools

Many metagenomics or metatranscriptomics analysis tools were conceived for the analysis of bacterial communities. For example, CLARK [16, 17] is a tool for the taxonomic classification of metagenomic reads using known bacterial genomes. GOTTCHA [18] is a taxonomic profiler that uses nonredundant signature databases for prokaryotic and viral genomes. Genometa [19] is a Java program to identify bacterial species and gene content from high-throughput datasets. MetaPhyler [20] estimates bacterial composition from metagenomic samples.

Others use a subset of the sequences for taxonomic profiling of metagenomes. Web-based solutions are provided by MG-RAST [21] and EBI metagenomics [22] that automatically analyse rRNA and mRNA in submitted samples. MetaPhlAn2 [23] and mOTU [24] use a subset of marker genes for taxonomic profiling. QIIME [25] uses Operational Taxonomic Units (OTUs) to assign a taxonomy.

A user-specified library of genomes of species that are present in the samples has to be provided for recent programs utilizing k-mers such as Kraken [26], LMAT [27] or DUDes [28].

The last category of tools searches the NCBI database to assign reads to taxonomical level after a BLAST search, including MEGAN [29] and Taxator-tk [30] or after a mapping with Bowtie, e.g. Centrifuge [31].

Four our purposes, we found that each existing tool exhibited a shortcoming that rendered it unsuitable for the read-level assignment of taxonomic and functional information to microeukaryotic sequences. We summarize our requirements versus the properties of existing tools in Table 1.

**Table 1.**
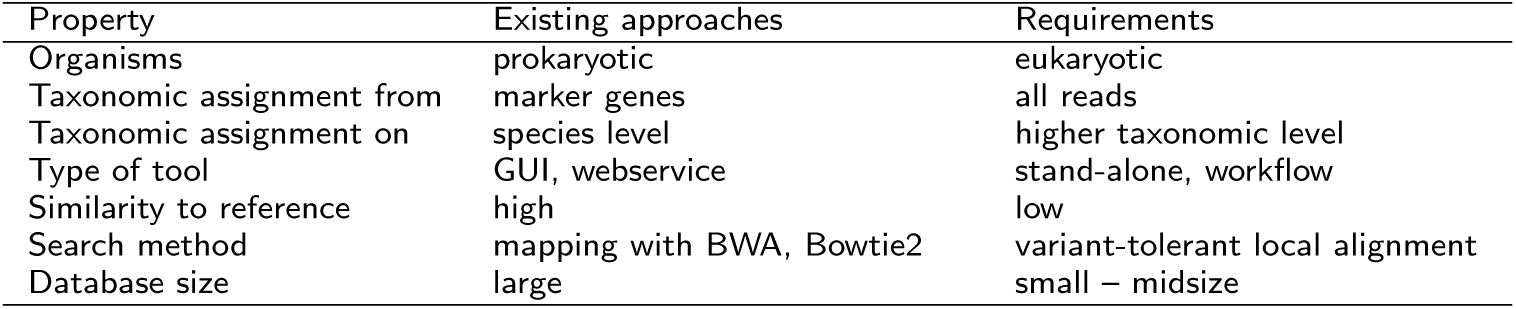
Issues with properties of existing approaches. Properties of existing approaches versus requirements for microeukaryotic environmental sequence analysis.

## Methodology and implementation

### Reference database

To counter-balance the uneven diversity of eukaryotic microorganisms present in public databases, we construct the TaxMapper reference database such that it evenly includes genomic and transcriptomic sequences from all eukaryotic supergroups and taxonomic groups. References from the databases of NCBI [32] and the Marine Microbial Eukaryote Transcriptome Sequencing Project [1] were selected based on the following criteria: (i) The taxa represent the main lineages within each of the seven supergroups of eukaryotes (see Fig. 1). (ii) Their genomes or transcriptomes are mostly complete; i.e., we excluded obviously incomplete datasets that consisted of only some hundred sequences. We thus selected 142 transcriptomes and genomes; the selection is described under “Results”.

**Figure 1.**
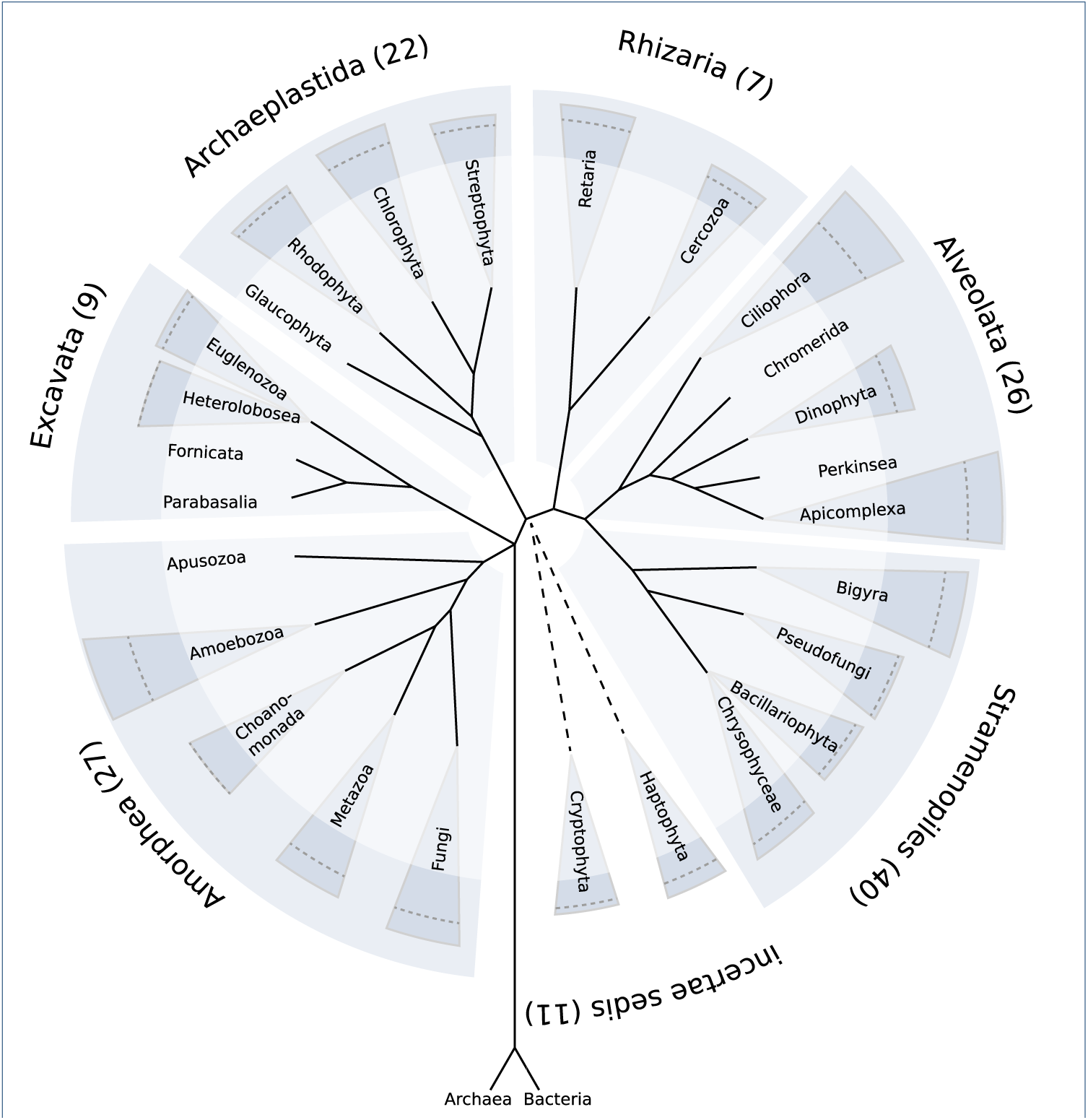
Taxonomy of eukaryotes. Taxonomy of eukaryotes with the supergroups and groups used in the reference database. Two remaining groups combining small lineages are not depicted. Coloured with darker background is the diversity of the supergroups and groups computed as the maximum Bray-Curtis dissimilarity over 4-mer spectra from the proteins of the reference genomes, as defined in [55]. Additionally, the mean Bray-Curtis dissimilarity is indicated as a dashed line. The taxonomy is based on Boenigk and Wodniok [52].

The protein sequences of all reference genomes or transcriptomes were downloaded, redundant sequences were discarded for each species and the amino acid sequences were used to build a database index.

### TaxMapper

TaxMapper is designed to allow an easy-to-use search with sequence reads in the compiled database and to filter erroneous hits. It consists of five modules (search, map, filter, count, plot) that can be run individually with user defined parameters or as a single step with default settings.

The initial *search* in the indexed database is conducted for a single read file or forward and reverse reads in parallel using the protein similarity search tool RAPSearch2 [33] (v2.24, fast mode, using a loose E-value cutoff of 10^5^, but restricted to the best 20 hits). RAPSearch2 performs a fast similarity search in a reduced amino-acid search space. The best 20 hits are returned for each query (read) sequence and *mapped* to the 7 taxonomic supergroups and 28 main lineages. Two hits are kept subsequently, the best hit (BH) and the next best hit, according to E-value, that falls into another lineage (next lineage hit, NLH). (Hits that are better than the NLH and agree with the taxonomic group of the BH are skipped.) Forward and reverse results can be combined by choosing either the option “best” to use the better of both searches or “concordant”, where forward and reverse have to map to the same taxon.

The *filter* idea behind TaxMapper is to assign taxonomic information only if the BH and NLH are “different enough”. If the differences between BH and NLH in mapping properties such as the E-value, identity, alignment score etc. are large, the assignment of the best hit is regarded trustworthy and is returned, otherwise no taxonomic group is ascribed to reduce false positive assignments. The details of the filter approach are discussed below (Subsection Filtering). Fig. 2 illustrates the difference of this approach to other approaches that use only the best hit or the lowest common ancestor (LCA) of several hits. While the best hit approach returns just the best hit, regardless of further results that might be equally good, the lowest common ancestor approach returns the lowest level in the taxonomic tree that the hits have in common, which might be close to the root if the hits are too diverse.

**Figure 2.**
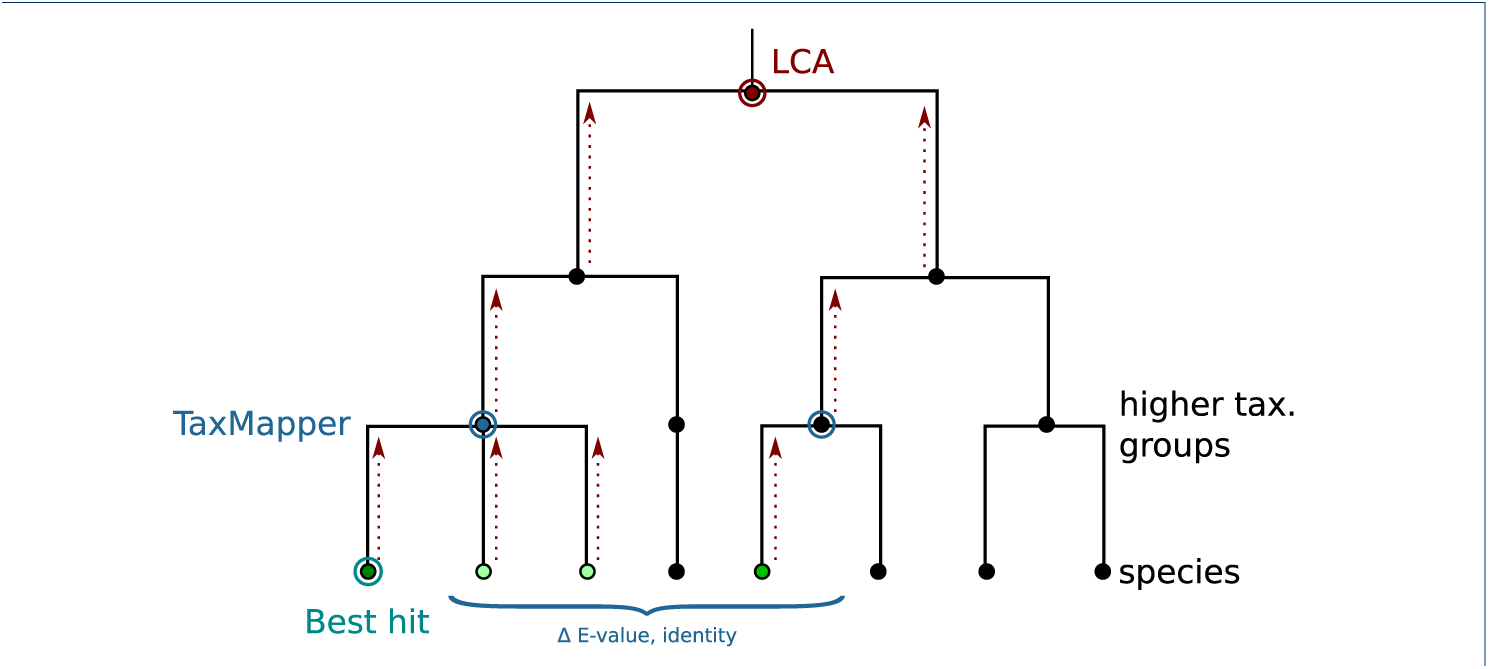
Differences between TaxMapper, LCA and best hit. Given the green leaves as possible hits, with the best hit circled in green, TaxMapper compares the best hits on a higher taxonomic level (blue circle) and uses the better hit (blue node) if the differences between the hits are large enough, while LCA is a bottom-up method that possibly returns the root of the taxonomy (red node) if the hits are too diverse.

Subsequently, *count* matrices can be generated over samples, summarizing the reads for all taxonomic groups to apply total count normalization and *plot* community compositions.

TaxMapper is implemented as a stand-alone tool in the Python language (v3.5). The statistical model for the filtering step (described below) was estimated using the generalized linear model function in R, applying maximum likelihood estimation (MLE). R is not required for running the TaxMapper software. TaxMapper can be run either stepwise with user-defined settings or for easier handling in one analysis step with default parameters. In the second case, just a folder of raw data in FASTQ or FASTA format has to be provided and all results are generated automatically. The analysis can be parallelized by declaring the number of threads to use and it is suggested to run it on a multicore machine or server for large datasets.

### Filtering

The filtering step based on the best hit (BH) and the nearest lineage hit (NHL) is a distinguishing feature of TaxMapper. Since we found it impossible to separate correct from incorrect taxonomic assignments based on BH and NLH E-values alone, we estimated a logistic regression model based on five BH/NLH properties:

1. percent identity of the BH,
2. ratio of identities between BH and NLH,
3. log_10_ E-value of BH,
4. difference in log_10_ E-values of BH and NLH,
5. the total size (in basepairs) in the database of the BH’s taxonomic group

The base frequencies were added as an independent variable in addition to the mapping statistics (E-value and identity) to include the different number of sequences per taxonomic group, which can bias hits toward more abundant taxa in the reference database.

In general, the binary logistic model is used to estimate the probability of a binary response *y* ∈ {0,1}, based on one or more independent variables (*x*_1_,…,*x_p_*):

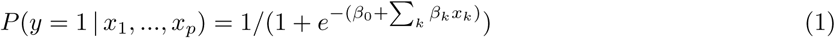

Here the *x_k_* are the five hit properties described above, and *y* = 1 corresponds to the event that the BH is a correct assignment, whereas *y* = 0 means that the BH is an incorrect assignment. The goal is to search for values of the coefficients *β* such that the probability *P*(*y* = 1 | *x*) is large when the hit properties *x* indicate that BH and NLH are sufficiently different such that the taxonomic assignment based on the BH is correct.

For estimating and testing the classifier, reads were chosen from 18 species that are included in the reference database and 17 species that are not included in the database, but where the taxonomic lineage is known and present in the database. Not all of the 28 groups could be used, since for some groups all available species were included in the database and further species for testing were not obtainable.

We obtained raw read data belonging to the above 35 species, listed with accession number in the supplementary file Suppl_TestTable.csv. Since for these reads, we know the correct taxonomic origin, we sorted them into two classes based on TaxMapper’s best hit (BH) alone: correctly classified or misclassified. We randomly chose 500 000 correctly classified (true positive, TP) and 500 000 misclassified (false positive, FP) reads as training data for estimating the model (see Fig. 3). This dataset of one million reads was split into 20% holdout data and 80% training and test data. The training and test data was again randomly split into 80% training and 20% test data 100 times to train and evaluate the classifier using 100-fold Monte Carlo cross-validation. In addition, in each cross validation round, the holdout data and randomly created reads were used to evaluate the classifier. Performance on the random reads (which by definition have no relation to any database sequence) allows us to estimate how well we are able to reject sequences that are from none of the eukaryotic lineages contained in the database. Results are given in the “Results” section.

**Figure 3.**
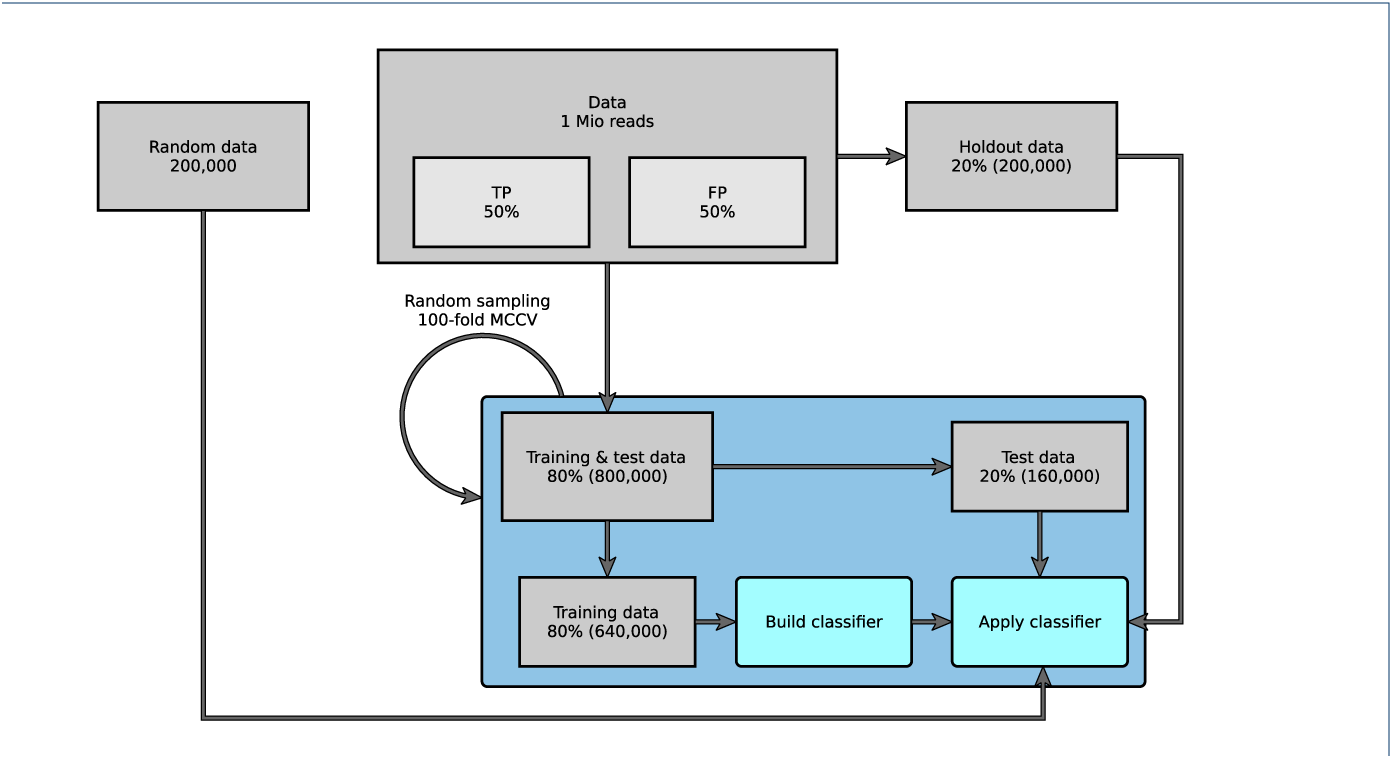
Classification scheme. One million reads from different taxonomic groups with 50% false positive and 50% true positive best hit assignments were used. This dataset was split in 20% holdout data and 80% training and test data, of which again 80% were used to train and 20% to test the classifier applying 100-fold Monte Carlo cross-validation. In addition, in each fold the holdout data and randomly simulated (nonsense) reads were used to evaluate the classifier.

## Workflow

A comprehensive workflow for metatranscriptome analysis was developed and made available in an executable Snakemake-based workflow. Snakemake is a workflow description language and execution environment developed by Köster *et al.* [34]. The workflow steps are defined in terms of rules with input, output and Shell, Python or R code. Dependencies between rules are automatically resolved and rules are automatically parallelized where possible. It features an easy to read, self-documenting syntax which also serves for version and parameter tracking. For the described workflow Snakemake version 3.9.1 was used.

The workflow covers both taxonomic assignment of each read (using TaxMapper) and functional assignment (using RAPSearch2 on the UniProt database). Steps and parameters can be adjusted using a provided configuration file (`config.yaml`).

In the following, the most important rules and steps of the workflow are explained. An overview is given in Figure 4.

**Figure 4.**
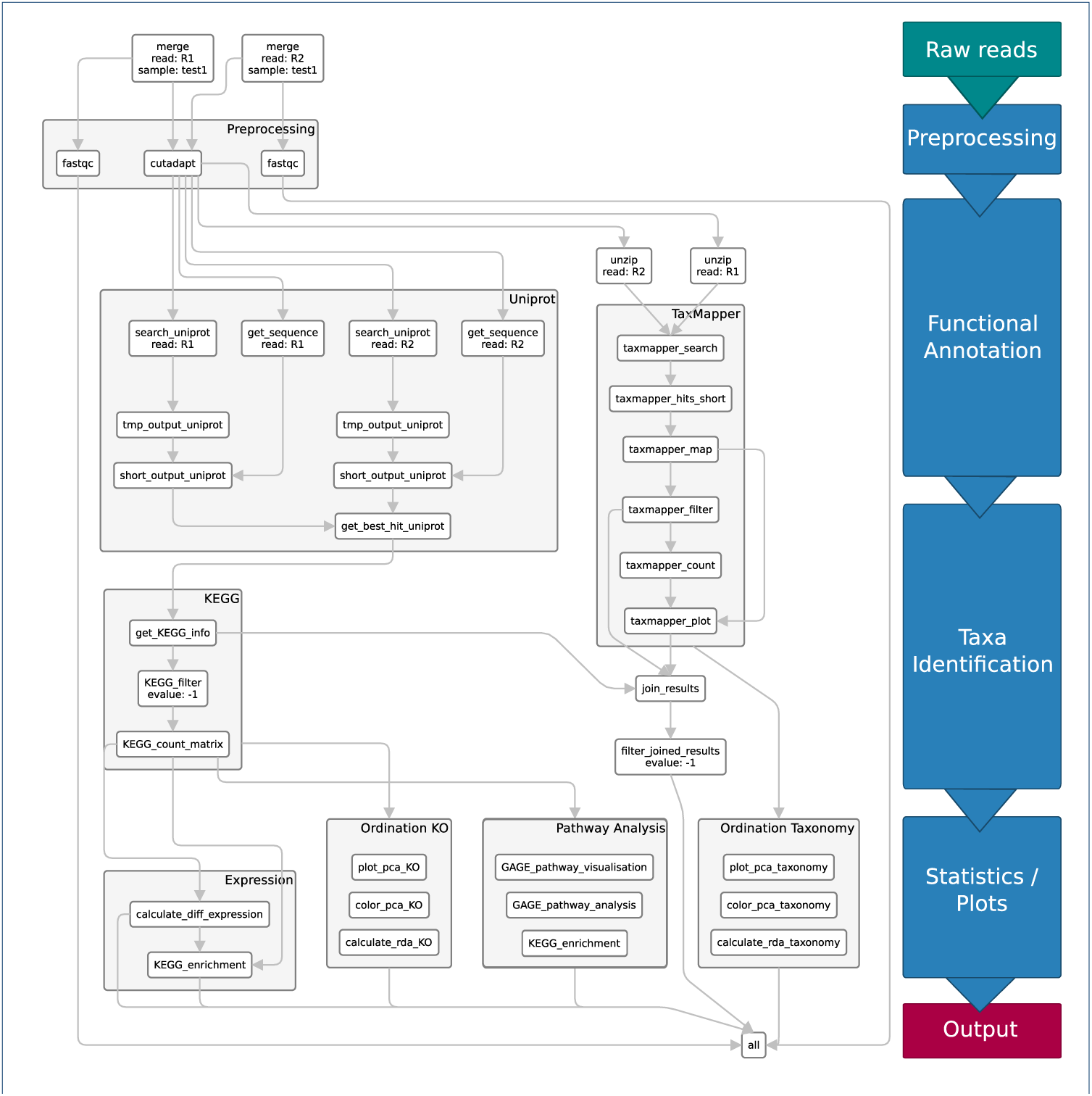
Snakemake workflow. Rules of the Snakemake workflow processing the FASTQ input files to the final output *all.*

The steps of the bioinformatic workflow are specified in the workflow management system Snakemake. Snakemake rules describe how to create output files from input files by executing commands on the input files. The commands can also be run on single files in the terminal, Python or R, but for automation, parallelization and reproducibility of the workflow, Snakemake is used. We briefly explain the Snakemake syntax here on a short exemplary Snakemake file:

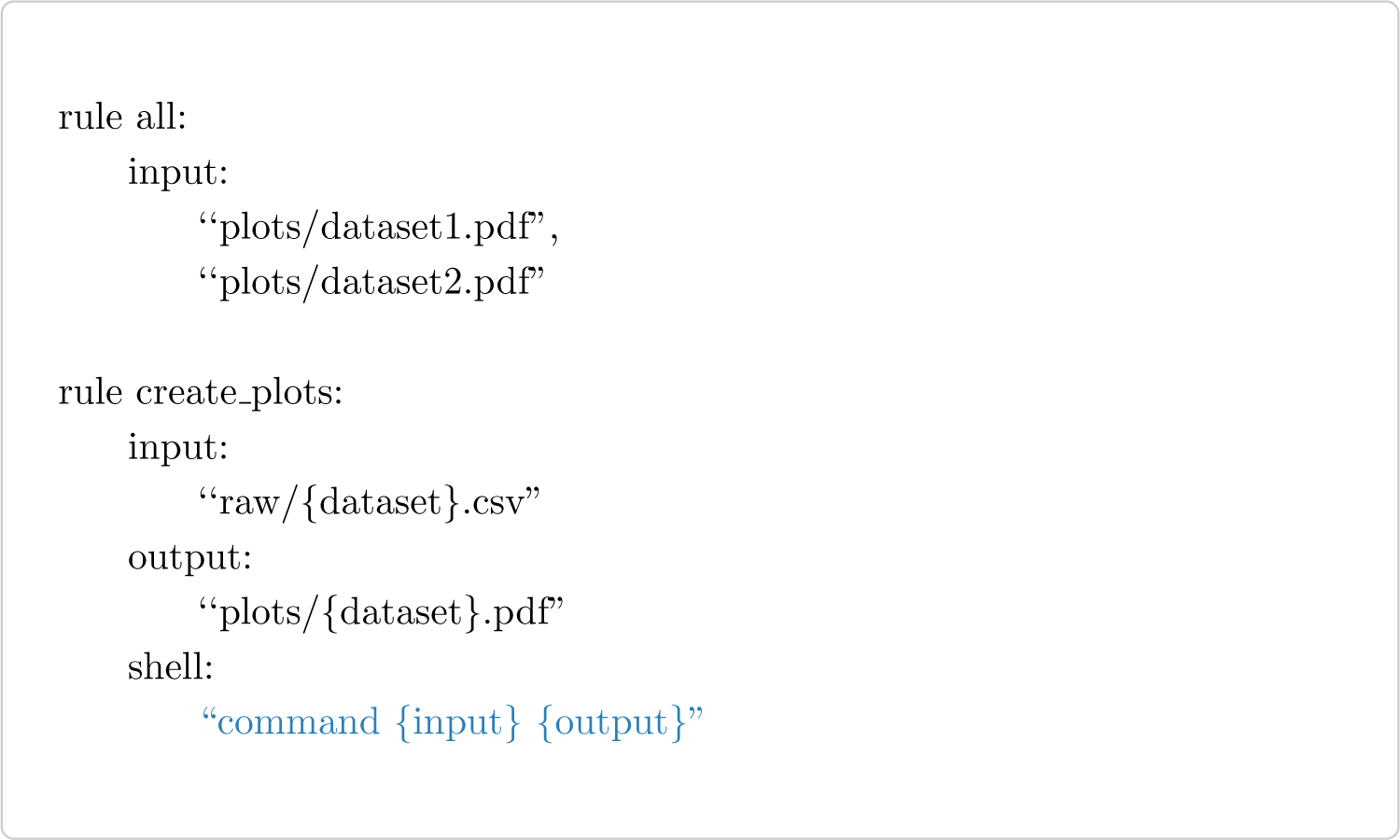

The desired final outputs of the workflow are described in the rule all, these are “plots/dataset1.pdf” and “plots/dataset2.pdf”. To create the plot, we run a shell command in the rule create_plots on the input “raw/{`dataset`}.csv” to create the output “plots/{`dataset`}.pdf”. Snakemake determines the rule dependencies by matching file names and automatically fills the wildcard `dataset` with the correct names: datasetl and dataset2, that are expected as the input of rule all.

### Preprocessing

The quality of raw sequencing reads is analysed using the quality control tool FastQC [35]. It computes various quality measures such as the base quality, overrepresented sequences, read length et cetera. The compressed FASTQ files are used as input and the snakemake rule runs FastQC as a shell command on the input. The wildcards `sample` and `pair` represent the sample name and forward and reverse read respectively.

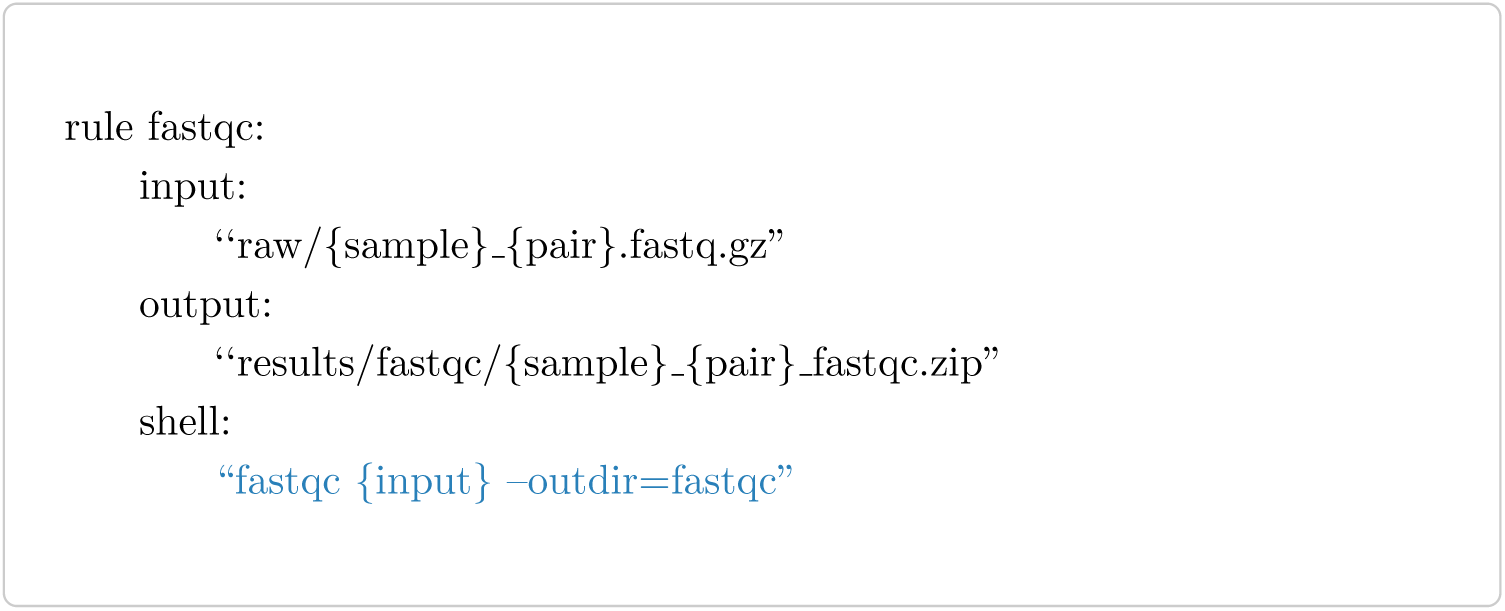

Identified low quality bases and sequencing adapters can be removed with trimming tools such as cutadapt (vl.12, [36]). From the forward and reverse read, given as input, the adapter beginning with ‘GATCGGAAGAGCA’ and bases with a quality value below 20 are trimmed. If the remaining read length is below 50, the whole read will be discarded. All output files are saved in the folder results/cleaned.

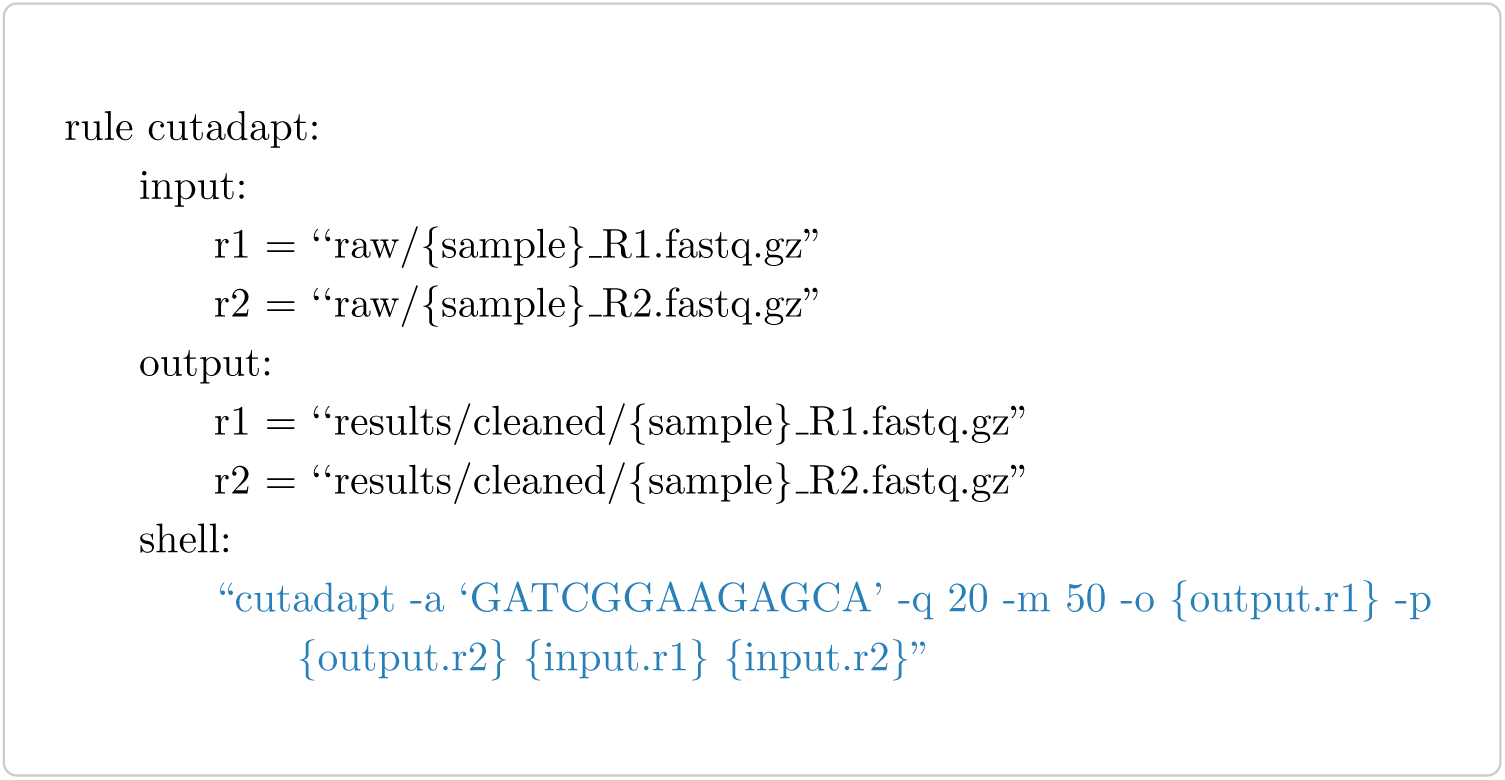

### Taxa identification

TaxMapper is used for the assignment and filtering of taxonomic information. For brevity, the one-step version is shown below, since it just needs an input folder with all FASTQ files and parallelization is performed within TaxMapper (here 20 threads are used via option `–t`). We have to get the input folder from the input files and provide an output file from TaxMapper as output for snakemake. The expand command is used to get a list of all input files by filling in the wildcards for `sample` and `pair`, which are lists of all filenames and forward and reverse reads provided in the configuration file. The database index is created within the subworkflow `taxonomy` which is given as the input database. To let Snakemake handle parallelization and provide user-defined parameters, the workflow can also be run in five successive steps: search, map, filter, count and plot (see Fig. 4 TaxMapper box).

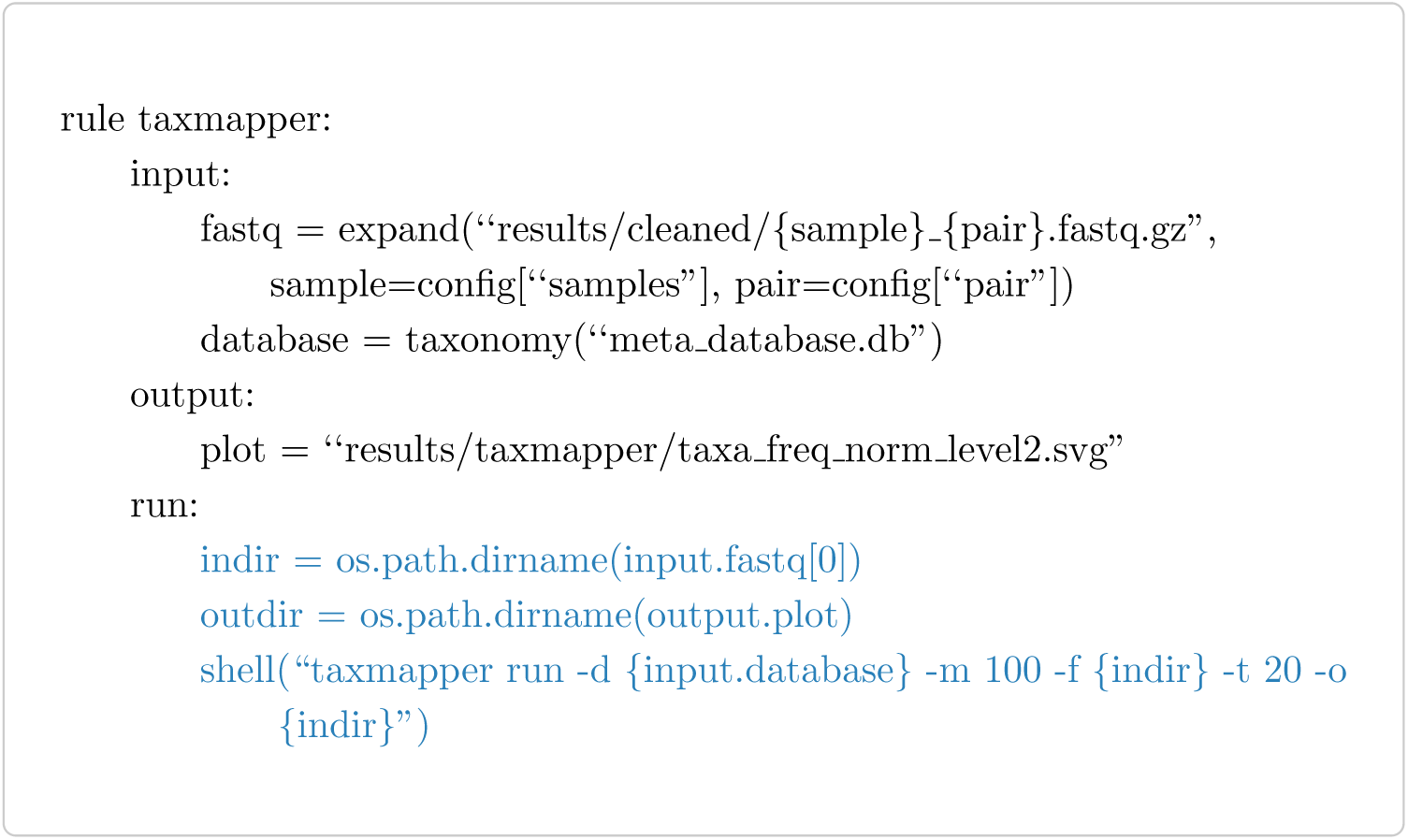

### Functional annotation

RAPSearch (v2.24, [33]), a fast protein similarity search tool, is used to search the read sequences in the Uniprot database (release 2016_06) [37]. The Uniprot database is downloaded and indexed as part of the workflow (in a subworkflow termed uniprot). The similarity search is performed with default parameters and the best hit is returned. Via a Uniprot identifier mapping file, obtained from the Uniprot database, KEGG (Kyoto Encyclopedia of Genes and Genomes, [38]) Orthology identifiers can be assigned to the query sequence.

Additional rules are used to shorten the output and combine the forward and reverse read mapping (see Fig. 4 Uniprot box). The input FASTQ files have to be first extracted from the gz archive to use them as input for RAPSearch2, then they are searched against Uniprot returning the alignments of the best hit or no result for each read.

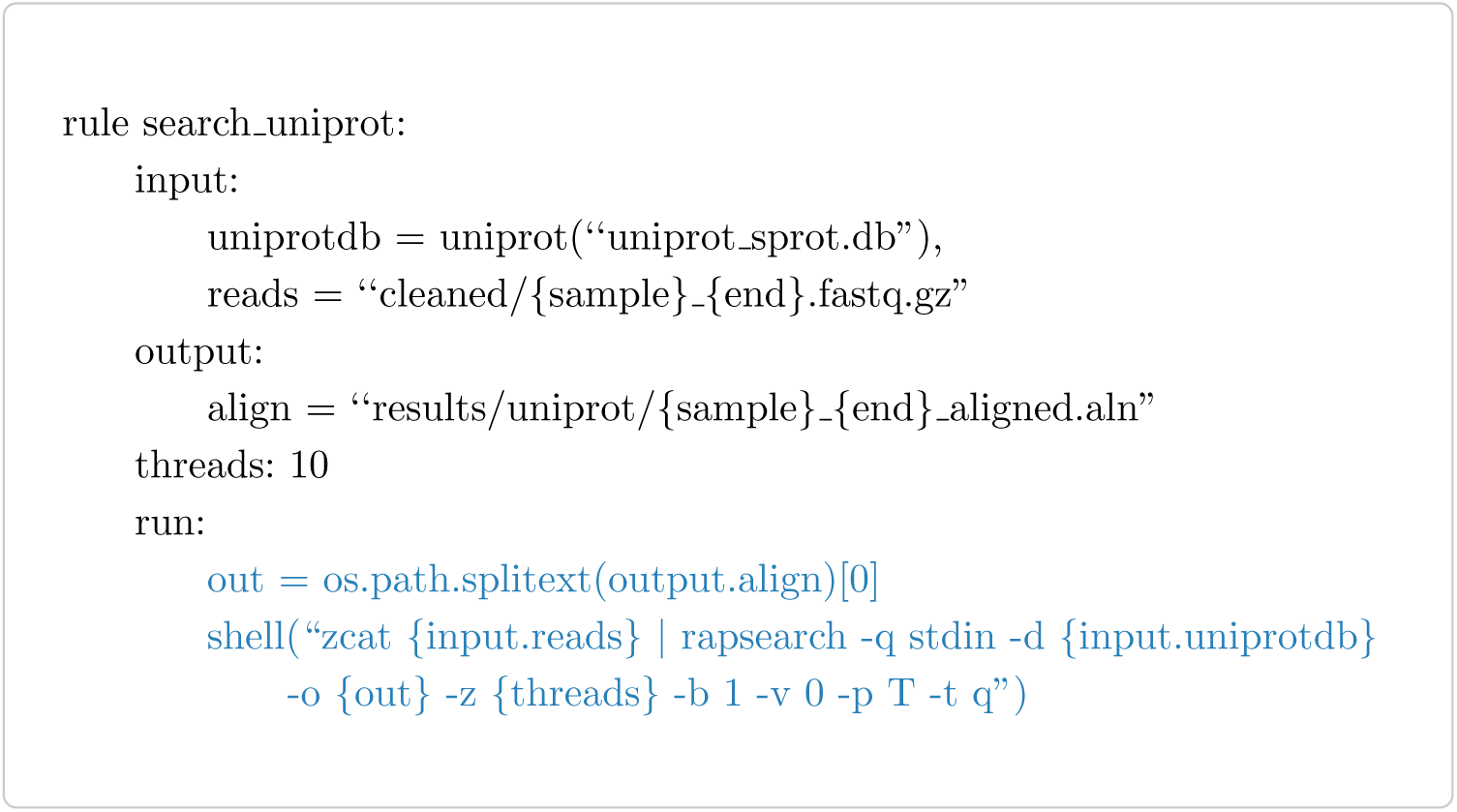

### Statistics and downstream analysis

Subsequent statistical analyses depend on the type of study and question. Since it is not always possible or intended to perform e.g. differential expression analysis, we included several possible rules in the workflow. All of the rules execute R code that is longer than a couple of lines and therefore not depicted here.

Existing rules include a differential expression analysis given different conditions using the Bioconductor package edgeR (v3.14.0, [39]), ordination analyses such as principal component analysis and redundancy analysis using the R package vegan (v2.3-4, [40]) and KEGG pathway analyses with the R packages GAGE (v2.21.1, [41]) and pathview (v1.9.0, [42]).

## Results and discussion

### Reference database

According to our criteria, 142 reference sequences were selected for the TaxMapper reference database (for details see Suppl file: Suppl_TaxTable.csv). These references belong to the seven supergroups of eukaryotes, including 28 main lineages. In accordance with the taxonomy published by Boenigk and Wodniok [43] and with the tree of life project [44], we chose different levels of each lineage to cover their molecular and functional diversity. Figure 1 and Table 2 give an overview.

**Table 2.**
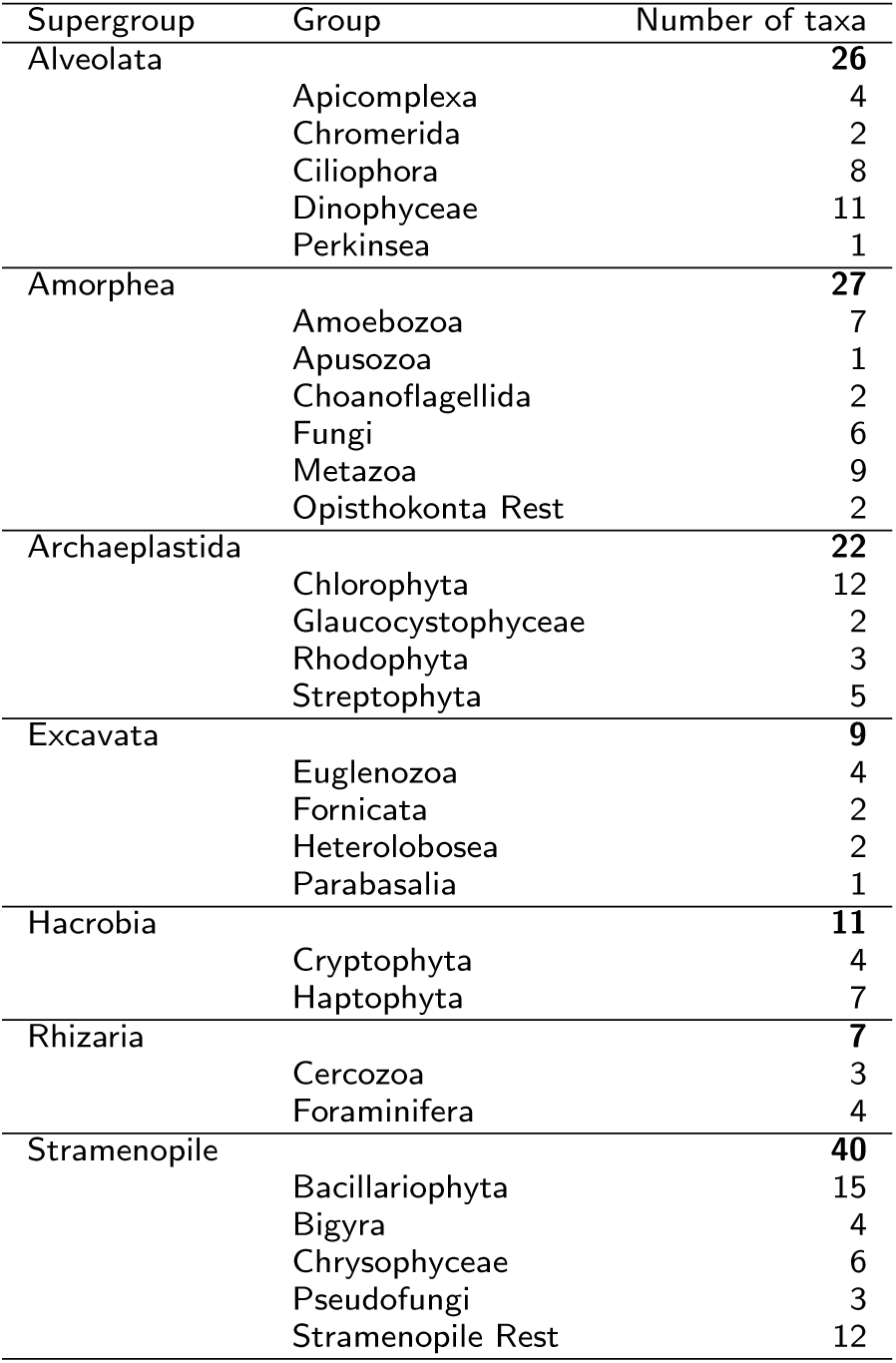
Number of taxa in used taxonomic groups. Bold numbers: number of taxa used for each supergroup; non-bold: number of taxa used for each taxonomic group in the reference database.

The supergroup Amorphea consists of two main lineages, the Opisthokonta (Holomycota and Holozoa) and Amoebozoa. Additionally, the small phylum Apusozoa is considered as a likely paraphyletic sistergroup of the Opistokonta [45, 46]. In the database the Amorphea are represented by 27 reference taxa. 19 taxa are affiliated with the Opisthokonta, including fungi representing the Holomycota, and Eumetazoa, Choanoflagellida (Choanomonada) and basal Opisthokonta, e.g. Filastera and Ichthyosporea here called Opisthokonta Rest, as representatives for the Holozoa. The Amoebozoa contain 7 reference taxa including lobose Amoebae, Archamoebae and Mycetozoa (slime moulds). One reference taxa is included for the phylum Apusozoa.

The supergroup Excavata is a very diverse group that can be summarized into two main groups, the Discoba including the lineages Euglenozoa, Heterolobosea and Jakobida as well as the Metamonada including the lineages Parabasalia and Fornicata. Many species of this supergroup are parasites [5] but some taxa e.g. most Euglenida are free-living and often occur in freshwater [47]. In the database the Excavata are represented by 9 reference taxa affiliated with Euglenozoa, Heterolobosea, Parabasalia and Fornicata. Due to few available transcriptomes of this supergroup in public databases and the focus on free-living taxa, only few references could be added.

The supergroup Archaeplastida includes three main lineages, the species-poor Glaucophyta (Glaucocystophyceae), the mostly marine Rhodophyta and the species-rich Viridiplantae (Chlorophyta, Streptophyta). Particularly the Chlorophyta are important primary producers in freshwater habitats [48]. Therefore, Archaeplastida are represented by 22 reference taxa affiliated with Chlorophyta, Streptophyta, Rhodophyta and Glaucocystophyceae.

The supergroup Rhizaria is a diverse group and consists of two main lineages, Cercozoa and Retaria (Foraminifera and Radiolaria). Cercozoa are very abundant in soil but can also occur in freshwaters and marine habitats [49]. In the database Rhizaria are represented by only 7 taxa belonging to Cercozoa and Foraminifera as there are only a few sequenced species available in public databases, particularly from Cercozoa.

The supergroup Alveolata is a very diverse group. It consists of three main lineages, Ciliophora, Apicomplexa and Dinophyta. Further, the smaller lineages Chromerida, Colpodellids and Perkinsea are affiliated with the Alveolata. Ciliophora and Dinophyceae can occur in high abundances and are important predators of other protists [50, 51]. Due to their importance and diversity they are covered by a high number of reference taxa (26) in the database: Ciliophora, Apicomplexa, Dinophyceae, Chromerida and Perkinsea.

The supergroup Stramenopiles is a very diverse group including many lineages which can be summarized into three groups, the Pseudofungi, the heterotrophic Bigyra and the plastid bearing Ochrophyta [52]. Some of these lineages, e.g. Bacillariophyta and Chrysophyceae, are very abundant in freshwater habitats [48, 50]. They are important primary producers and predators of bacteria. Therefore, we covered this group by a high number of 40 reference taxa. Pseudofungi were included as well as Bigyra summarizing the three lineages Bicosoecida, Blastocystis and Labyrinthulida. The Ochrophyta are represented by the two abundant freshwater groups Bacillariophyta and Chrysophyceae and a collection of other reference taxa affiliated with several Stramenopile lineages called Stramenopiles Rest.

An additional “group” in the eukaryotic tree of life are the incertae sedis Eukaryota which include amongst others the Hacrobia (Cryptophyta, Haptophyta) [5]. The evolutionary position of theses taxa is still uncertain as the phylogenetic position differs depending on the studied organism and genes. In the database Hacrobia are represented by 11 reference taxa, affiliated with Cryptophyta and Haptophyta.

### Evaluation of the filtering step

After training the classifier to reject assignments of training reads whose best hit misses the correct taxonomic group, we evaluated the performance on the test, random and holdout dataset.

The results are depicted as receiver operating characteristic (ROC) curves in Fig. 5 A and compared based on the area under the curve (AUC) and accuracy (ACC) in Table 3. Shown are true positive rate (TPR) and false positive rate (FPR) of TaxMapper results varying over the cutoff for the probability *P*(*y* = 1|*x*_1_,…, *x*_5_). Results are also given when no logistic model, but a simple E-value cutoff for the best hit, is used.

**Figure 5.**
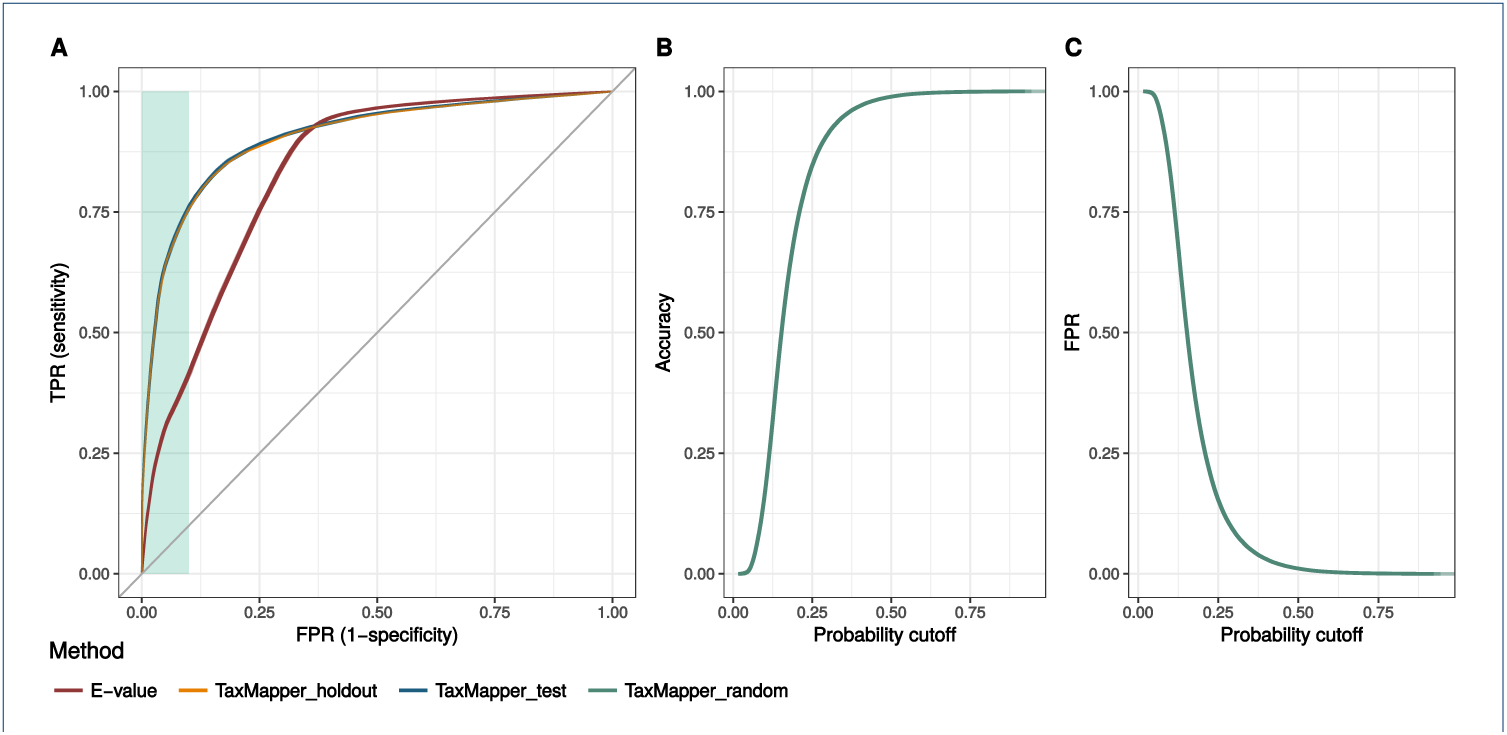
Validation. (A) False positive rate (FPR, x-axis) versus true positive rate (TPR, y-axis) of TaxMapper results on test dataset (blue), TaxMapper result on holdout data (orange), and simple E-value cutoff results on test dataset (red). (The blue and orange curves overlap in the subfigure on the left side.) The green background indicates the desired area with a low FPR (≤ 0.1). (B) Accuracy of TaxMapper on the random nonsense data (green) against the probability cutoff. (C) False positive rate of TaxMapper on the random data over all probability cutoffs.

**Table 3.**
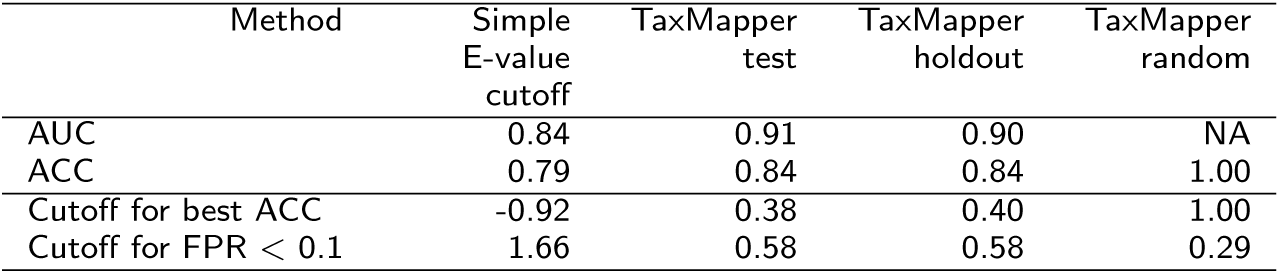
Evaluation of TaxMapper. Comparison of area under the ROC curve (AUC) and accuracy (ACC) for the E-value cutoff (test data) and TaxMapper on test, holdout and random data. The cutoffs leading to the best results in ACC and a false positive rate below 0.1 are shown below.

| Method | Simple E-value cutoff | TaxMapper test | TaxMapper holdout | TaxMapper random |
| --- | --- | --- | --- | --- |
| AUC | 0.84 | 0.91 | 0.90 | NA |
| ACC | 0.79 | 0.84 | 0.84 | 1.00 |
| Cutoff for best ACC | -0.92 | 0.38 | 0.40 | 1.00 |
| Cutoff for FPR < 0.1 | 1.66 | 0.58 | 0.58 | 0.29 |

TaxMapper yields superior results, especially in the desired area with low false positive rates, and an AUC of 0.90–0.91 in contrast to 0.84 for the simple E-value cutoff method. The highest accuracy of 0.84 was obtained for a probability cutoff of 0.38 and 0.40 for TaxMapper (test and holdout data, respectively). The best accuracy (0.79) for a simple E-value cutoff lay below –0.92 (log_10_ E-value).

A false positive rate below 0.1 could be obtained with a probability cutoff of 0.58 or log_10_ E-value below 1.66. Obviously, in the random dataset only the number of false positives can be reduced, resulting in the best accuracy of 1.0 for a probability cutoff of 1.0, filtering out all reads. But as shown in Fig. 5 B and C, the accuracy increases rapidly and a low false positive rate below 0.1 is already obtained with an average probability cutoff of 0.29 (see Fig. 5 and Tab. 3).

### Evaluation of TaxMapper against other tools

The runtime and results of TaxMapper were compared to the tool Taxator-tk and Centrifuge, to our knowledge the only tools that can be run on a server and assign sequences to a taxonomy on read-level (see Fig. 6). Both tools were run with default parameters and as described in the manual. As a reference for Centrifuge the non redundant NCBI index was used as provided by the authors of Centrifuge. For Taxator-tk the provided refpacks could not be used, since they focus on prokaryotic taxa, therefore a refpack using the NCBI nr database was build according to the instructions on the website. The search step of Taxator-tk utilises a blastn or LAST search against the NCBI nonredundant nucleotide database. Due to the long runtime, only the holdout data with 200 000 reads was tested. Overall, Taxator-tk using the Megan algorithm takes 3980:13 minutes, Centrifuge takes 15:07 minutes and TaxMapper 32:49 minutes (wall clock time) on a server with AMD Opteron processors (6176, 2.3 GHz) using 20 threads. This corresponds to a user time of 182:18 minutes for TaxMapper, of which the search step takes longest with 180:23 minutes.

**Figure 6.**
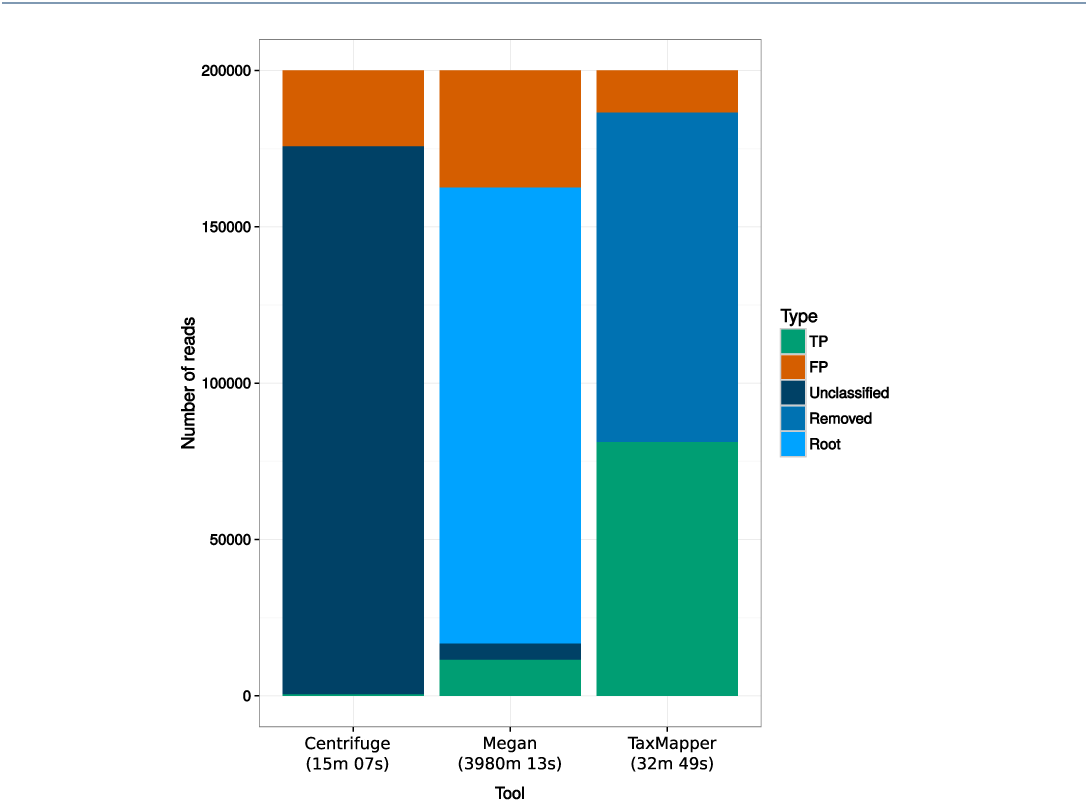
Comparison to other tools. Shown are the results obtained on the holdout dataset using the tool Centrifuge, TaxMapper and Taxator-tk with the Megan algorithm and the required wall clock time in brackets (run with 20 threads in parallel). Depicted are the number of reads resulting in a true positive (TP) assignment, false positive (FP) assignment, unclassified taxonomy, reads mapping to the root of the taxonomic tree and filtered (removed) reads by TaxMapper.

Centrifuge uses the fast mapping algorithm Bowtie to map the reads against the NCBI database. The drawback is that Bowtie allows few mismatches and therefore reads map only to very similar sequences. If the organism or a close relative is not contained in the database, a taxonomy cannot be assigned, leading to many unclassified reads for this method. The Megan algorithm of Taxator-tk uses BLAST, therefore only few reads are unclassified, but the majority map to the root node of the taxonomy, due to the lowest common ancestor approach described in Figure 2. The original algorithm developed for Taxator-tk is optimized for longer reads, starting with 500 bp, and was not used here. TaxMapper results in the highest number of true positive assignments and the lowest number of false positives. Results were the taxonomic assignment of the best hit was insecure, were removed in the filter step.

## Discussion

### Example application: sliver dataset

To showcase an application, the metatransciptome workflow was run on a subset of sequencing data from a study published in 2014 by Boenigk *et al.* [52]. In brief, a short-term silver exposure experiment was conducted on nine 20 L plastic tanks containing water from a natural plankton community from an eutrophic pond at the campus Essen of the University Duisburg-Essen. The nine tanks were divided into three experimental groups (control, silver nitrate and silver nanoparticle exposure) with three replicate tanks each. The subsample used here contains the control samples and the silver nitrate samples. The metatranscriptomic workflow was applied to analyse the functional and taxonomic differences between the treatments. Figure 7 A depicts the community compositions with the largest changes visible in the groups Bacillariophyta and Chlorophyta. The taxonomic changes are also depicted in the PCA in Figure 7 B, separating on the second principal component the control samples from the samples treated with a sublethal silver concentration of 5 μg/L. On the functional level a test for differential expression reveals 34 KEGG orthologous genes that differ significantly (FDR < 0.1) between the two groups and show an enrichment of photosynthesis pathways. It is known that silver ions affect the primary metabolism in particular photosynthesis by direct interference [52, 53]. On the other hand, it has been shown that for low concentrations of silver green algae grows is increased as observed in Figure 7 A [54].

**Figure 7.**
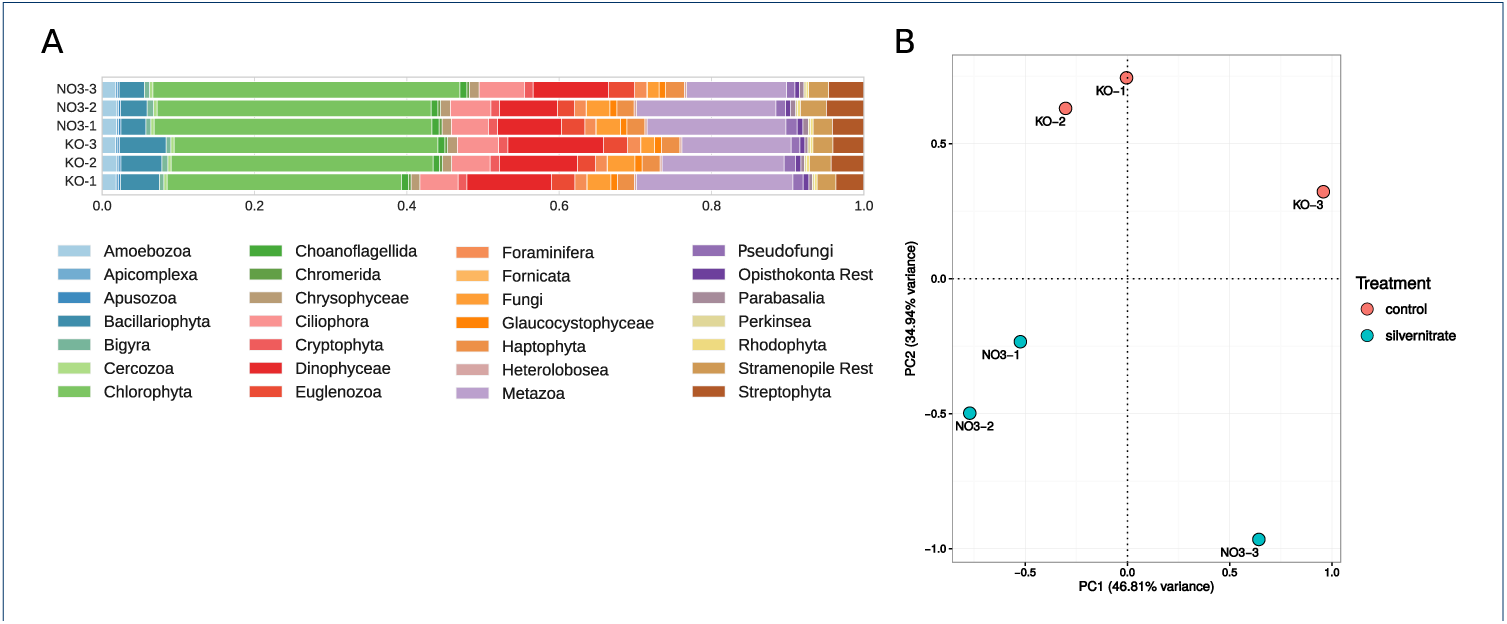
Community composition and principal component analysis of silver dataset. A) community composition of a subset of data from a metatranscriptome sequencing study, where the effect of silver nitrate was tested on the community and function. B) principal component analysis (PCA) of the TMM-normalized taxonomic count data, colored according to treatment: control in red and samples with added silver nitrate in blue.

A subset of this study with the first 100,000 reads per FASTQ file is provided with the workflow as test dataset.

### Future database updates

When new sequences become available which further complete the diversity of the eukaryotic supergroups, an update of the database will be released. In particular, the Excavata and Rhizaria should be extended in future versions, for which at the moment only few appropriate genomes or transcriptomes are present.

## Conclusions

Despite the large number of tools developed for taxonomic analyses, the majority of them aims at different sequencing data (e.g. rRNA, contigs) or organismic groups (prokaryotes) and does not allow a combined functional and taxonomic analysis of metatranscriptomic data. We therefore developed the presented tool TaxMapper to work in conjunction with a constructed microeukaryotic reference database for taxonomic assignment, and included the taxonomic analysis in a complete workflow for metatranscriptomic sequence analysis.

The smaller, but more appropriate reference for protists, allows a faster search than a comparable search against whole NCBI.

False positive assignments can be filtered using a probability cutoff on a logistic regression model based on features of the best hit and next lineage hit, which yielded better result than a simple E-value cutoff.

TaxMapper can be run straightforwardly on a folder of sequencing data or as part of the Snakemake workflow. The workflow performs quality assessment, functional and taxonomic annotation and (multivariate) statistical analyses using available environmental factors or different sample groups. The provided workflow ensures a reproducible analysis which can be easily extended to new samples.

Both the TaxMapper tool and the workflow are available as open-source software at Bitbucket under the MIT license: https://bitbucket.org/dbeisser/taxmapper and as a Bioconda package: https://bioconda.github.io/recipes/taxmapper/README.html.

## Declarations

### Ethics approval and consent to participate

Not applicable.

### Consent to publish

Not applicable.

### Availability of data and material

The data and software are available at Bitbucket https://bitbucket.org/dbeisser/taxmapper, https://bitbucket.org/dbeisser/taxmapper_supplement and as a Bioconda package: https://bioconda.github.io/recipes/taxmapper/README.html.

### Author’s contributions

DB compiled the reference database, developed the tool TaxMapper and the Snake-make workflow and wrote the manuscript. NG selected the taxa for the reference database and wrote the Reference Database section in the manuscript. LG provided the first version of reference taxa. HT contributed to the tool TaxMapper, created the conda package and performed code review. JB provided expertise on protists and the eukaryotic phylogeny. SR provided bioinformatics expertise and wrote sections of the manuscript. JB and SR led and guided the study. All authors participated in writing and approved the final version of the manuscript.

## Acknowledgements

We thank Matthias Höller for running Centrifuge on the holdout data.

## List of abbreviations

ACC: Accuracy
AUC: Area under the curve
BH: Best hit
DFG: Deutsche Forschungsgemeinschaft
FDR: False discovery rate
FP: False postive
FPR: False positive rate
HTS: High-throughput sequencing
KEGG: Kyoto Encyclopedia of Genes and Genomes
LCA: Lowest common ancestor
MLE: Maximum likelihood estimation
MMETSP: Marine Microbial Eukaryote Transcriptome Sequencing Project
NCBI: National Center for Biotechnology Information
NLH: Next lineage hit
OTU: Operational Taxonomic Unit
PCA: Principal component analysis
ROC: Receiver operating characteristic
TMM: Trimmed mean of M-values
TP: True positive
TPR: True positive rate

## Additional files

Suppl_TaxTable.csv — Taxa contained in reference database

Information on taxa contained in reference database, including taxonomic affiliation, accession number and database.

Suppl_TestTable.csv — Validation taxa

Information on taxa used for evaluating the logistic regression model.

## References

1. Keeling, P.J., Burki, F., Wilcox, H.M., Allam, B., Allen, E.E., Amaral-Zettler, L.A., Armbrust, E.V., Archibald, J.M., Bharti, A.K., Bell, C.J., Beszteri, B., Bidle, K.D., Cameron, C.T., Campbell, L., Caron, D.A., Cattolico, R.A., Collier, J.L., Coyne, K., Davy, S.K., Deschamps, P., Dyhrman, S.T., Edvardsen, B., Gates, R.D., Gobler, C. J., Greenwood, S.J., Guida, S.M., Jacobi, J.L., Jakobsen, K.S., James, E.R., Jenkins, B., John, U., Johnson, M.D., Juhl, A.R., Kamp, A., Katz, L.A., Kiene, R., Kudryavtsev, A., Leander, B.S., Lin, S., Lovejoy, C., Lynn, D., Marchetti, A., McManus, G., Nedelcu, A.M., Menden-Deuer, S., Miceli, C., Mock, T., Montresor, M., Moran, M.A., Murray, S., Nadathur, G., Nagai, S., Ngam, P.B., Palenik, B., Pawlowski, J., Petroni, G., Piganeau, G., Posewitz, M.C., Rengefors, K., Romano, G., Rumpho, M.E., Rynearson, T., Schilling, K.B., Schroeder, D.C., Simpson, A.G.B., Slamovits, C.H., Smith, D.R., Smith, G.J., Smith, S.R., Sosik, H.M., Stief, P., Theriot, E., Twary, S.N., Umale, P.E., Vaulot, D., Wawrik, B., Wheeler, G.L., Wilson, W.H., Xu, Y., Zingone, A., Worden, A.Z.: The Marine Microbial Eukaryote Transcriptome Sequencing Project (MMETSP): Illuminating the Functional Diversity of Eukaryotic Life in the Oceans through Transcriptome Sequencing. PLoS Biology 12(6), 1001889 (2014). doi:10.1371/journal.pbio.1001889

2. de Vargas, C., Audic, S., Henry, N., Decelle, J., Mahe, F., Logares, R., Lara, E., Berney, C., Le Bescot, N., Probert, I., Carmichael, M., Poulain, J., Romac, S., Colin, S., Aury, J.-M., Bittner, L., Chaffron, S., Dunthorn, M., Engelen, S., Flegontova, O., Guidi, L., Horak, A., Jaillon, O., Lima-Mendez, G., Luke, J., Malviya, S., Morard, R., Mulot, M., Scalco, E., Siano, R., Vincent, F., Zingone, A., Dimier, C., Picheral, M., Searson, S., Kandels-Lewis, S., Acinas, S.G., Bork, P., Bowler, C., Gorsky, G., Grimsley, N., Hingamp, P., Iudicone, D., Not, F., Ogata, H., Pesant, S., Raes, J., Sieracki, M.E., Speich, S., Stemmann, L., Sunagawa, S., Weissenbach, J., Wincker, P., Karsenti, E., Boss, E., Follows, M., Karp-Boss, L., Krzic, U., Reynaud, E.G., Sardet, C., Sullivan, M.B., Velayoudon, D.: Eukaryotic plankton diversity in the sunlit ocean. Science 348(6237), 1261605–1261605 (2015). doi:10.1126/science.1261605

3. Escobar-Zepeda, A., Vera-Ponce de León, A., Sanchez-Flores, A.: The Road to Metagenomics: From Microbiology to DNA Sequencing Technologies and Bioinformatics. Frontiers in genetics 6(DEC), 348 (2015). doi:10.3389/fgene.2015.00348

4. Schlegel, M., Hülsmann, N.: Protists – A textbook example for a paraphyletic taxon. Organisms Diversity & Evolution 7(2), 166–172 (2007). doi:10.1016/j.ode.2006.11.001

5. Burki, F.: The Eukaryotic Tree of Life from a Global Phylogenomic Perspective. Cold Spring Harbor Perspectives in Biology 6(5), 016147–016147 (2014). doi:10.1101/cshperspect.a016147

6. Grossmann, L., Jensen, M., Heider, D., Jost, S., Glüucksman, E., Hartikainen, H., Mahamdallie, S.S., Gardner, M., Hoffmann, D., Bass, D., Boenigk, J.: Protistan community analysis: key findings of a large-scale molecular sampling. The ISME Journal 10(9), 2269–2279 (2016). doi:10.1038/ismej.2016.10

7. Finlay, B.J., Esteban, G.F.: Freshwater protozoa: biodiversity and ecological function. Biodiversity & Conservation 7(9), 1163–1186 (1998). doi:10.1023/A:1008879616066

8. Ackermann, B., Esser, M., Scherwaß, A., Arndt, H.: Long-Term Dynamics of Microbial Biofilm Communities of the River Rhine with Special References to Ciliates. International Review of Hydrobiology 96(1), 1–19 (2011). doi:10.1002/iroh.201011286

9. Geisen, S., Tveit, A.T., Clark, I.M., Richter, A., Svenning, M.M., Bonkowski, M., Urich, T.: Metatranscriptomic census of active protists in soils. The ISME Journal 9(10), 2178–2190 (2015). doi:10.1038/ismej.2015.30

10. Flynn, K.J., Stoecker, D.K., Mitra, A., Raven, J.A., Glibert, P.M., Hansen, P.J., Granéli, E., Burkholder, J.M.: Misuse of the phytoplankton-zooplankton dichotomy: The need to assign organisms as mixotrophs within plankton functional types. Journal of Plankton Research 35(1), 3–11 (2013). doi:10.1093/plankt/fbs062

11. Šimek, K., Hartman, P., Nedoma, J., Pernthaler, J., Springmann, D., Vrba, J., Psenner, R.: Community structure, picoplankton grazing and Zooplankton control of heterotrophic nanoflagellates in a eutrophic reservoir during the summer phytoplankton maximum. Aquatic Microbial Ecology 12(1), 49–63 (1997). doi:10.3354/ame012049

12. Mitra, A., Flynn, K.J., Burkholder, J.M., Berge, T., Calbet, A., Raven, J.A., Granéli, E., Glibert, P.M., Hansen, P.J., Stoecker, D.K., Thingstad, F., Tillmann, U., Våge, S., Wilken, S., Zubkov, M.V.: The role of mixotrophic protists in the biological carbon pump. Biogeosciences 11(4), 995–1005 (2014). doi:10.5194/bg-11-995-2014

13. Leimena, M.M., Ramiro-Garcia, J., Davids, M., van den Bogert, B., Smidt, H., Smid, E.J., Boekhorst, J., Zoetendal, E.G., Schaap, P.J., Kleerebezem, M.: A comprehensive metatranscriptome analysis pipeline and its validation using human small intestine microbiota datasets. BMC Genomics 14, 530 (2013). doi:10.1186/1471-2164-14-530

14. Goncalves, A., Tikhonov, A., Brazma, A., Kapushesky, M.: A pipeline for RNA-seq data processing and quality assessment. Bioinformatics 27(6), 867–869 (2011). doi:10.1093/bioinformatics/btr012

15. Marchetti, a., Schruth, D.M., Durkin, C.a., Parker, M.S., Kodner, R.B., Berthiaume, C.T., Morales, R., Allen, a.E., Armbrust, E.V.: Comparative metatranscriptomics identifies molecular bases for the physiological responses of phytoplankton to varying iron availability. Proceedings of the National Academy of Sciences 109(6), 317–325 (2012). doi:10.1073/pnas.1118408109

16. Ounit, R., Wanamaker, S., Close, T.J., Lonardi, S.: CLARK: fast and accurate classification of metagenomic and genomic sequences using discriminative k-mers. BMC Genomics 16(1), 236 (2015). doi:10.1186/s12864-015-1419-2

17. Ounit, R., Lonardi, S.: Higher classification sensitivity of short metagenomic reads with CLARKS. Bioinformatics 32(24), 3823–3825 (2016). doi:10.1093/bioinformatics/btw542

18. Freitas, T.A.K., Li, P.-E., Scholz, M.B., Chain, P.S.G.: Accurate read-based metagenome characterization using a hierarchical suite of unique signatures. Nucleic Acids Research 43(10), 180 (2015). doi:10.1093/nar/gkv180

19. Davenport, C.F., Neugebauer, J., Beckmann, N., Friedrich, B., Kameri, B., Kokott, S., Paetow, M., Siekmann, B., Wieding-Drewes, M., Wienhöfer, M., Wolf, S., Tümmler, B., Ahlers, V., Sprengel, F.: Genometa - A Fast and Accurate Classifier for Short Metagenomic Shotgun Reads. PLoS ONE 7(8), 41224 (2012). doi:10.1371/journal.pone.0041224

20. Liu, B., Gibbons, T., Ghodsi, M., Pop, M.: Metaphyler: Taxonomic profiling for metagenomic sequences. In: 2010 IEEE International Conference on Bioinformatics and Biomedicine (BIBM), pp. 95–100 (2010). doi:10.1109/BIBM.2010.5706544

21. Meyer, F., Paarmann, D., D’Souza, M., Olson, R., Glass, E., Kubal, M., Paczian, T., Rodriguez, A., Stevens, R., Wilke, A., Wilkening, J., Edwards, R.: The metagenomics RAST server – a public resource for the automatic phylogenetic and functional analysis of metagenomes. BMC Bioinformatics 9(1), 386 (2008). doi:10.1186/1471-2105-9-386

22. Mitchell, A., Bucchini, F., Cochrane, G., Denise, H., ten Hoopen, P., Fraser, M., Pesseat, S., Potter, S., Scheremetjew, M., Sterk, P., Finn, R.D.: EBI metagenomics in 2016 - an expanding and evolving resource for the analysis and archiving of metagenomic data. Nucleic Acids Research 44(D1), 595–603 (2016). doi:10.1093/nar/gkv1195

23. Segata, N., Waldron, L., Ballarini, A., Narasimhan, V., Jousson, O., Huttenhower, C.: Metagenomic microbial community profiling using unique clade-specific marker genes. Nature Methods 9(8), 811–814 (2012). doi:10.1038/nmeth.2066

24. Sunagawa, S., Mende, D.R., Zeller, G., Izquierdo-Carrasco, F., Berger, S.a., Kultima, J.R., Coelho, L.P., Arumugam, M., Tap, J., Nielsen, H.B., Rasmussen, S., Brunak, S., Pedersen, O., Guarner, F., de Vos, W.M., Wang, J., Li, J., Doré, J., Ehrlich, S.D., Stamatakis, A., Bork, P.: Metagenomic species profiling using universal phylogenetic marker genes. Nature Methods 10(12), 1196–1199 (2013). doi:10.1038/nmeth.2693

25. Caporaso, J.G., Kuczynski, J., Stombaugh, J., Bittinger, K., Bushman, F.D., Costello, E.K., Fierer, N., Peña, A.G., Goodrich, J.K., Gordon, J.I., Huttley, G.a., Kelley, S.T., Knights, D., Koenig, J.E., Ley, R.E., Lozupone, C. a., McDonald, D., Muegge, B.D., Pirrung, M., Reeder, J., Sevinsky, J.R., Turnbaugh, P.J., Walters, W.a., Widmann, J., Yatsunenko, T., Zaneveld, J., Knight, R.: QIIME allows analysis of high-throughput community sequencing data. Nature Methods 7(5), 335–336 (2010). doi:10.1038/nmeth.f.303

26. Wood, D.E., Salzberg, S.L.: Kraken: ultrafast metagenomic sequence classification using exact alignments. Genome Biology 15(3), 46 (2014). doi:10.1186/gb-2014-15-3-r46

27. Ames, S.K., Hysom, D.A., Gardner, S.N., Lloyd, G.S., Gokhale, M.B., Allen, J.E.: Scalable metagenomic taxonomy classification using a reference genome database. Bioinformatics 29(18), 2253–2260 (2013). doi:10.1093/bioinformatics/btt389

28. Piro, V.C., Lindner, M.S., Renard, B.Y.: DUDes: A top-down taxonomic profiler for metagenomics. Bioinformatics 32(15), 2272–2280 (2016). doi:10.1093/bioinformatics/btw150

29. Huson, D.H., Auch, A.F., Qi, J., Schuster, S.C.: MEGAN analysis of metagenomic data. Genome Research 17(3), 377–386 (2007). doi:10.1101/gr.5969107

30. Dröge, J., Gregor, I., McHardy, A.C.: Taxator-tk: precise taxonomic assignment of metagenomes by fast approximation of evolutionary neighborhoods. Bioinformatics 31(6), 817–824 (2015). doi:10.1093/bioinformatics/btu745

31. Kim, D., Song, L., Breitwieser, F.P., Salzberg, S.L.: Centrifuge: rapid and sensitive classification of metagenomic sequences. Genome Research 26(12), 1721–1729 (2016). doi:10.1101/gr.210641.116

32. Coordinators, N.R.: Database Resources of the National Center for Biotechnology Information. Nucleic Acids Research 45(D1), 12–17 (2017). doi:10.1093/nar/gkw1071

33. Zhao, Y., Tang, H., Ye, Y.: RAPSearch2: a fast and memory-efficient protein similarity search tool for next-generation sequencing data. Bioinformatics 28(1), 125–126 (2012). doi:10.1093/bioinformatics/btr595

34. Köster, J., Rahmann, S.: Snakemake–a scalable bioinformatics workflow engine. Bioinformatics 28(19), 2520–2522 (2012). doi:10.1093/bioinformatics/bts480

35. Andrews, S.: FastQC a quality control tool for high throughput sequence data

36. Martin, M.: Cutadapt removes adapter sequences from high-throughput sequencing reads. EMBnet.journal 17(1), 10–12 (2011). doi:10.14806/ej.17.1.200

37. Magrane, M., Consortium, U.: UniProt Knowledgebase: a hub of integrated protein data. Database 2011, 009–009 (2011). doi:10.1093/database/bar009

38. Kanehisa, M.: KEGG: Kyoto Encyclopedia of Genes and Genomes. Nucleic Acids Research 28(1), 27–30 (2000). doi:10.1093/nar/28.1.27

39. Robinson, M.D., McCarthy, D.J., Smyth, G.K.: edgeR: A Bioconductor package for differential expression analysis of digital gene expression data. Bioinformatics 26(1), 139–140 (2009). doi:10.1093/bioinformatics/btp616

40. Oksanen, J., Blanchet, F.G., Kindt, R., Legendre, P., Minchin, P.R., O’Hara, R.B., Simpson, G.L., Solymos, P., Stevens, M.H.H., Wagner, H.: vegan: Community Ecology Package (2016)

41. Luo, W., Friedman, M.S., Shedden, K., Hankenson, K.D., Woolf, P.J.: GAGE: generally applicable gene set enrichment for pathway analysis. BMC Bioinformatics 10(1), 161 (2009). doi:10.1186/1471-2105-10-161

42. Luo, W., Brouwer, C.: Pathview: an R/Bioconductor package for pathway-based data integration and visualization. Bioinformatics 29(14), 1830–1831 (2013). doi:10.1093/bioinformatics/btt285

43. Boenigk, J., Wodniok, S.: Biodiversität und Erdgeschichte. Springer, Berlin, Heidelberg (2014). doi:10.1007/978-3-642-55389-9

44. Maddison, D.R., Schultz, K.-S.: The Tree of Life Web Project. http://tolweb.org

45. Cavalier-Smith, T., Chao, E.E.: Phylogeny and Evolution of Apusomonadida (Protozoa: Apusozoa): New Genera and Species. Protist 161(4), 549–576 (2010). doi:10.1016/j.protis.2010.04.002

46. Paps, J., Medina-Chacón, L.A., Marshall, W., Suga, H., Ruiz-Trillo, I.: Molecular Phylogeny of Unikonts: New Insights into the Position of Apusomonads and Ancyromonads and the Internal Relationships of Opisthokonts. Protist 164(1), 2–12 (2013). doi:10.1016/j.protis.2012.09.002

47. Leander, B.S.: Euglenida (2012). http://tolweb.org/Euglenida/97461/2012.11.10

48. Garnier, J., Billen, G., Coste, M.: Seasonal succession of diatoms and Chlorophyceae in the drainage network of the Seine River: Observation and modeling. Limnology and Oceanography 40(4), 750–765 (1995). doi:10.4319/lo.1995.40.4.0750

49. Bass, D., Cavalier-Smith, T.: Cercozoa (2009). http://tolweb.org/Cercozoa/121187/2009.03.22

50. Auer, B., Arndt, H.: Taxonomic composition and biomass of heterotrophic flagellates in relation to lake trophy and season. Freshwater Biology 46(7), 959–972 (2001). doi:10.1046/j.1365-2427.2001.00730.x

51. Stoecker, D.K., Li, A.S., Coats, D.W., Gustafson, D.E., Nannen, M.K.: Mixotrophy in the dinoflagellate Prorocentrum minimum. Marine Ecology Progress Series 152(1-3), 1–12 (1997). doi:10.3354/meps152001

52. Boenigk, J., Beisser, D., Zimmermann, S., Bock, C., Jakobi, J., Grabner, D., Groβmann, L., Rahmann, S., Barcikowski, S., Sures, B.: Effects of silver nitrate and silver nanoparticles on a planktonic community: general trends after short-term exposure. PloS one 9(4), 95340 (2014). doi:10.1371/journal.pone.0095340

53. Beisser, D., Kaschani, F., Graupner, N., Grossmann, L., Jensen, M., Ninck, S., Schulz, S. Florian ANDandRahmann, Boenigk, J., Kaiser, M.: Quantitative proteomics reveals ecophysiological effects of light and silver stress on the mixotrophic protist poterioochromonas malhamensis. PLOS ONE 12(1), 1–20 (2017). doi:10.1371/journal.pone.0168183

54. Schmittschmitt, J.P., Shaw, J.R., Birge, W.J.: The 4th International Conference Proceedings: Transport, Fate and Effects of Silver in the Environment, pp. 245–249. University of Wisconsin System, Sea Grant Institute, Madison, WI (1996)

55. Dubinkina, V.B., Ischenko, D.S., Ulyantsev, V.I., Tyakht, A.V., Alexeev, D.G.: Assessment of k-mer spectrum applicability for metagenomic dissimilarity analysis. BMC Bioinformatics (2016). doi:10.1186/s12859-015-0875-7

